# A Theoretical Model of Telomere Length Regulation

**DOI:** 10.1101/602037

**Authors:** Haruhiko Ishii

## Abstract

Telomere length is maintained by a negative feedback mechanism that inhibits the telomerase activity when the telomere is sufficiently long. A prevailing explanation is that the negative feedback is caused by a mechanism that “counts” the number of regulatory protein molecules that are bound to the telomere. However, how such protein counting is accomplished is not clear. In this paper, I introduce a simple theoretical model to consider how the telomerase inhibition can depend on the telomere length. I show that this model is able to explain some key features of regulation of telomere length. While the real telomeres are more complex, this simple model may capture the essence of the mechanism.

## Introduction

A telomere is a structure found at each end of a linear chromosome (1). Telomeres usually have repetitive DNA sequences; they are repeats of TTAGGG in human and many other species. The repeats are added to the telomeres by the enzyme telomerase (2). The addition of the repeats counteracts the gradual loss of DNA that would otherwise occur due to inability of the DNA replication machinery to completely replicate the ends of linear DNA. The telomeres also protect ends of chromosomes from unwanted DNA repair responses that can lead to deleterious outcomes such as apoptosis, senescence, and chromosome fusions (3). The single-stranded 3’ overhang of the telomeric DNA invades the double-stranded repeats to form a lariat-like structure called a T-loop (4), instead of being exposed. The repeats also serve as binding sites for various telomeric proteins that play roles in protecting the telomere and regulating the length of the telomere (reviewed in (5–8)).

The lengths of telomeres are maintained in a range that is characteristic of the species and the cell type. Murray et al. (9) proposed that this telomere-length homeostasis is achieved by a negative feedback mechanism. According to this model, when a telomere is short, the telomerase can lengthen the telomere uninhibited. However, the telomerase activity is blocked when the telomere becomes longer. This leads to shortening of the telomere due to incomplete replication. A telomere approaches a steady state length by balancing of these processes. The finding that the telomerase preferentially adds repeats to short telomeres *in vivo* supports this model (10, 11).

The question is how this negative feedback is achieved. An attractive model is the protein-counting model (12), although other models have also been proposed (13, 14). As noted above, a number of proteins bind to the telomere, either directly to the DNA, or indirectly through protein-protein interactions. While the proteins that bind to the telomere differ considerably depending on the types of organisms (5–8), what is common is that many of them function as negative regulators of the telomere length: their mutations or RNAi knock down lead to lengthening of the telomeres; overexpressing them leads to shortening of the telomeres. The protein-counting model states that the negative feedback that inhibits the telomerase happens by sensing the number of the regulatory protein molecules that are bound to the telomere. Thus, the telomerase is inhibited when the telomere is long enough to accommodate a certain number of the protein molecules.

While the protein-counting model explains a number of experimental findings (12), how the “protein counting” occurs is not clear. A long telomere would naturally provide more binding sites for the proteins than a shorter one, but many of the binding sites would be far from the end of the telomere where the telomerase operates. It is not obvious how the proteins bound far from the telomerase can influence its activity.

In this paper, I propose a mechanism of how the occupancy of a telomere-binding protein can depend on the telomere length. I demonstrate this in a case of a simple theoretical model. The length-dependency is a consequence of assuming cooperative binding of the protein to the telomere. Even though the model described here is simplistic, the basic underlying mechanism may operate in more complex real telomeres.

## Results

### The basic premises of the model

I would like the model to be able to explain the following experimental observations:

1. Inhibition of the telomerase activity happens in a manner that depends on the length of a given telomere. The longer the telomere, more likely the telomerase activity is blocked.
2. Inhibition of the telomerase activity also depends on the abundance of the telomere-binding proteins. If the proteins are overexpressed, the telomerase activity is blocked at a shorter telomere length than in a normal condition. If the expression of the proteins is downregulated, the telomeres are extended longer than in a normal condition.

Many proteins associate with telomeres and they differ greatly depending on the organisms (5–8). If we try to account for all of the different proteins, the model will be very complicated and not easily tractable. The fact that different proteins in different organisms regulate telomere lengths in a similar manner suggests that there is a common mechanism that applies regardless of the specific details. What I would like to achieve is to build a simple conceptual model to capture the essence of the mechanism rather than building a detailed realistic model.

Here are several simplifications that I make.

1. In reality, there are many different proteins that bind to telomeres for a given species. However, I would like to represent the collective behavior of such telomere-binding proteins by one idealized protein P in the model.
2. I focus on modeling the T-loop. The size (the contour length) of the T-loop may not be identical to the total telomere length, but is constrained by the total telomere length. I will use the size of the T-loop as a proxy for the telomere length.
3. For simplicity, I idealize the T-loop as a homogeneous ring with binding sites for the protein P.
4. I assume that binding of P at the end of the telomere prevents the access of the telomerase. Since we are approximating the telomere as a homogeneous ring, the average binding of P per site represents the degree to which the telomerase is inhibited. I do not attempt to model lengthening and shortening of the telomere explicitly.

These simplifications reduce the problem to that of the protein P binding to a homogeneous one-dimensional lattice with a periodic boundary condition (Fig.1*A*). Each site can take one of two possible states: occupied, or unoccupied. I will use *s*_*j*_ to represent the state at the position *j*:

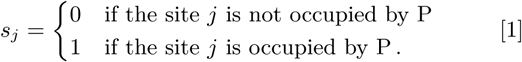

The total number of sites in this lattice *N* is a proxy for the length of the telomere that we are modeling. The state of the whole system can be represented by the vector {*s*_1_, *s*_2_, …, *s*_*N*_}.

**Fig. 1.**
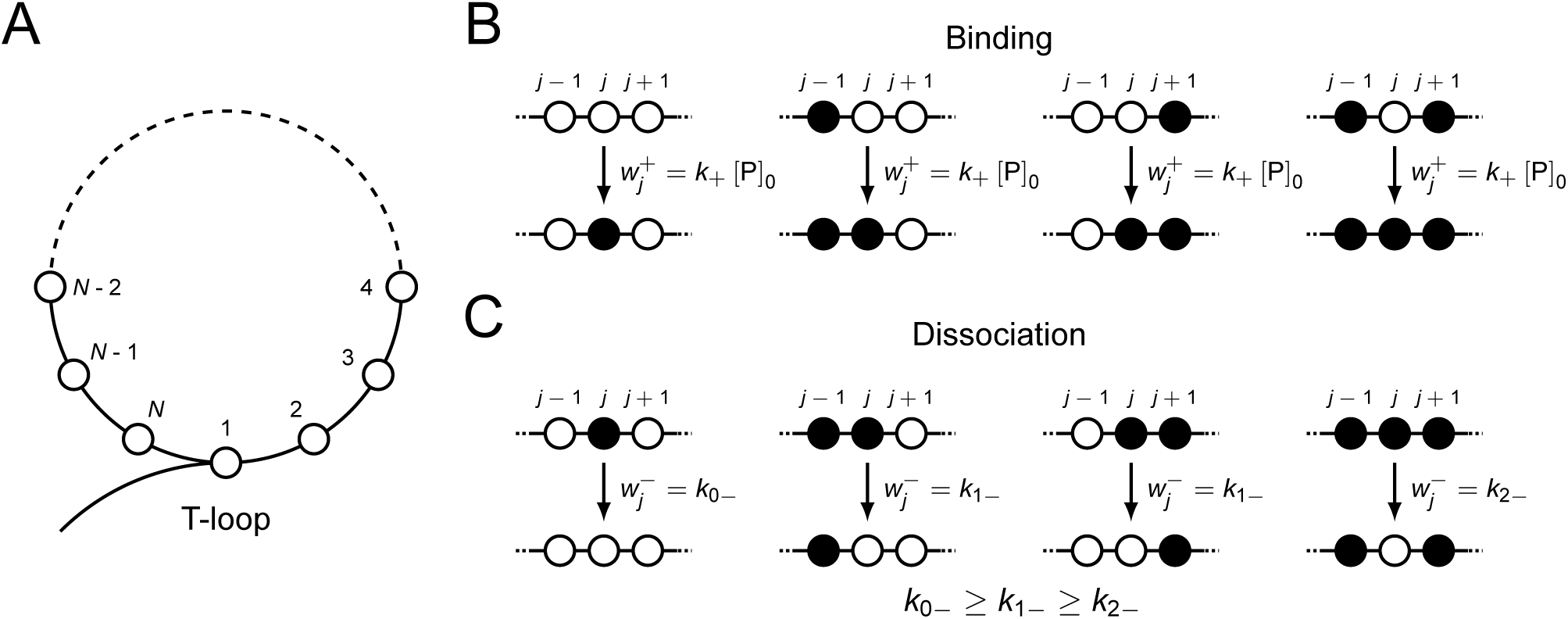
The model. (A) The T-loop is modeled as a ring with *N* binding sites. *N* gives a measure of the telomere size. (B) The open and filled circles represent unoccupied and occupied sites, respectively. The rate of binding of protein P to the site *j* is proportional to the concentration of free protein, [P]_0_, and does not depend on the states of the neighboring sites *j −* 1 and *j* + 1. (C) The rate of dissociation of protein P from the site *j* depends on the states of the neighboring sites *j −* 1 and *j* + 1. If *j −* 1 and *j* + 1 are unoccupied, the rate is *k*_0*−*_. If one of *j −* 1 or *j* + 1 is occupied, the rate is *k*_1*−*_. If both *j −* 1 and *j* + 1 are occupied, the rate is *k*_2*−*_.

### The dynamics

Suppose that the system evolves by stochastic binding and dissociation of P that changes the values of *s*_*j*_ for *j* = 1, 2,, *N*. I will use 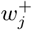 and 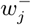 to represent the probabilities that *s*_*j*_ changes from 0 to 1 and 1 to 0 in a unit time, respectively.

Let us assume that the binding is driven by mass action —the probability of binding is proportional to the concentration of unbound P. If the probability of binding to site *j* does not depend on the occupancy of the neighboring sites, 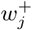 can be written in the following form:

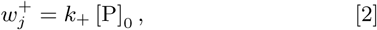

where [P]_0_ is the concentration of unbound P. (See Fig.1*B*.)

I would like to model that the rate of dissociation 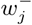 dependent on the occupancies of the neighboring sites because of the interactions between molecules of P bound to them. For simplicity, we will only consider the influence of the adjacent sites (Fig.1*C*):

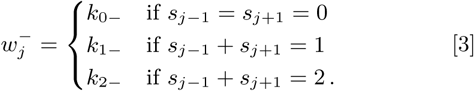

Assuming that binding of P to the site *j* is stabilized by other molecules occupying the neighboring sites,

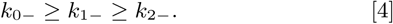

We can rewrite *k*_1*−*_ and and *k*_2*−*_ as

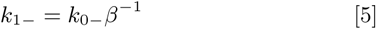

and

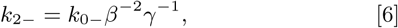

where *β* ≥ 1 and *βγ* ≥ 1. Then, 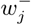 can be written as

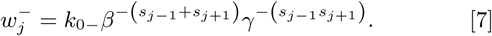

This model has similarity to the theoretical model of heterochromatin formation studied by Hodges and Crabtree (15). The difference is that in their model it is the binding rather than dissociation that is affected by the states of adjacent sites and the binding happens only by nucleation or propagation.

### The equilibrium properties

I would like to illustrate the properties of the model by calculating values when the system is in equilibrium. Let us use *p* (*s*_1_, *s*_2_,…, *s*_*N−*1_, *s*_*N*_; *t*) to represent the probability that the system is in the state {*s*_1_, *s*_2_, …,*s*_*N−*1_, *s*_*N*_} at time *t*. The rate that *s*_*j*_ switches from 0 to 1 at time *t* is 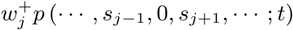. The rate that *s*_*j*_ switches from 1 to 0 at time *t* is 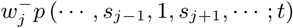. Consider a probability distribution that satisfies the following detailed balance:

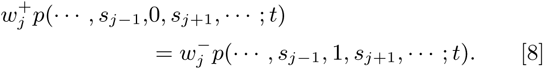

Such a distribution is in equilibrium and does not change with time because the rate of dissociation of the protein P from a given site *j* is always balanced by the rate of binding. I will represent the equilibrium distribution by *p*_eq_ (*s*_1_, *s*_2_, …, *s*_*N−*1_, *s*_*N*_).

The equilibrium distribution *p*_eq_ should satisfy the following relationship:

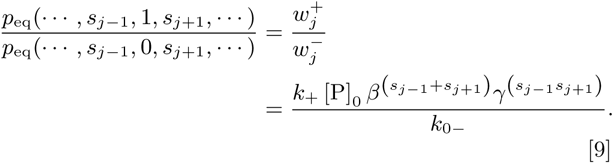

This becomes easier to picture by representing the probability distribution as the Boltzmann distribution with the following energy function *E*(*s*_1_, *s*_2_, …, *s*_*N*_) for the state {*s*_1_, *s*_2_, …, *s*_*N*_}:

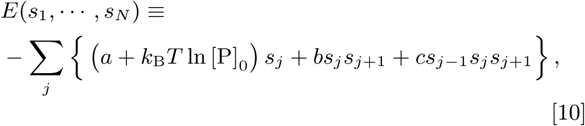

where we define *a, b*, and *c* as the following:

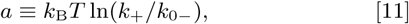

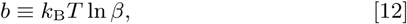

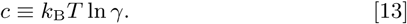

It is straightforward to see that the equilibrium probability *p*_eq_(…, *s*_*j−*1_, *s*_*j*_, *s*_*j*+1_, …) for any state {…, *s*_*j−*1_, *s*_*j*_, *s*_*j*+1_, …} is proportional to the Boltzmann weight:

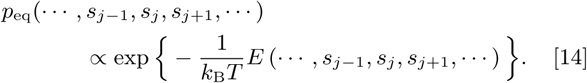

The way the stochastic system defined by the transition probabilities Eq.2 and Eq.7 leads to a thermal equilibrium described by Eq.14 is analogous to how systems driven by the Metropolis dynamics (16) or the Glauber dynamics (17) result in thermal equilibria. We can interpret the rates for binding and dissociation in terms of the Arrhenius theory—the activation energy for binding corresponds to

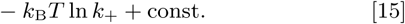

and the activation energy for dissociation from site *j* corresponds to

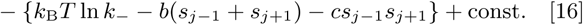

Solving the equilibrium properties of this model becomes an exercise of statistical mechanics. In particular, if *c* is equal to 0, this model is equivalent to the one-dimensional Ising model in a uniform magnetic field (18) just as in the case of the model of gene regulation that I studied previously (19). As long as *b* is non-zero, the model retains cooperativity. I will focus on the case where *c* is zero for simplicity.

I define the partition function as

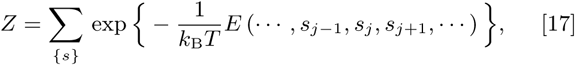

where the sum is for all possible combinations of {*s*_1_, *s*_2_, …, *s*_*N*_}.

Using the transfer matrix method (20), the partition function can be written as

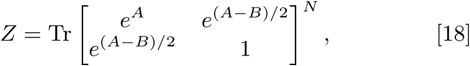

where

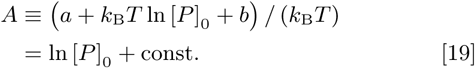

and

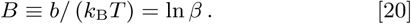

It is simple to diagonalize the matrix as

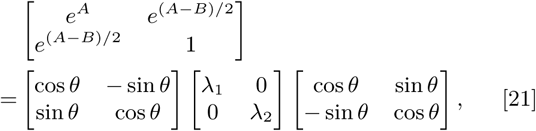

where

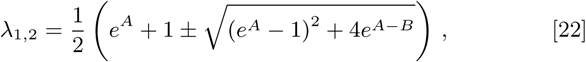

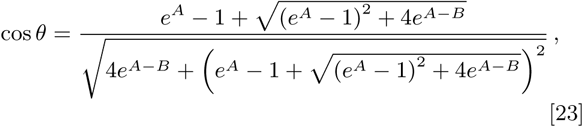

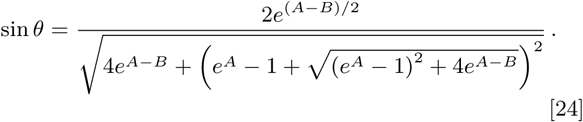

The partition function can be written using the eigenvalues *λ*_1,2_ as

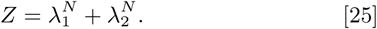

The expectancy that the site *j* is occupied is

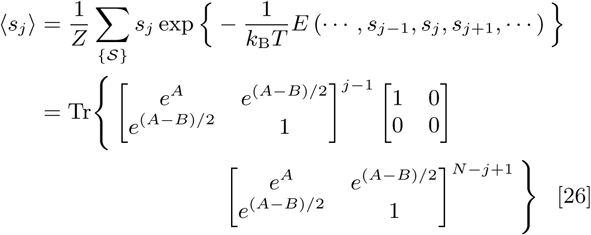

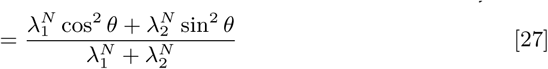

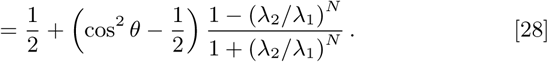

Note that, since we idealized the system as homogeneous, ⟨*s*_*j*_⟩ does not depend on the position *j*. Therefore, the occupancy at the very end of the telomere where the telomerase operates is the same as the occupancy anywhere else in this simplified model.

For large *N*, ⟨*s*_*j*_⟩ approaches

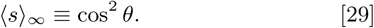

How fast ⟨*s*_*j*_⟩ approaches ⟨*s*⟩ _∞_ depends on *λ*_2_*/λ*_1_. We can introduce

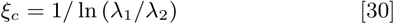

as the correlation length or the length scale. Then ⟨*s*_*j*_⟩ can be rewritten as

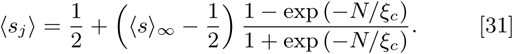

What this relationship shows is that ⟨*s*_*j*_⟩ does not reach the full occupancy ⟨*s*⟩_∞_ until *N* is roughly *ξ*_*c*_ or larger.

Fig.2(A), (B), and (C) show how ⟨*s*⟩_∞_ varies as a function *A* = ln [*P*]_0_ + const., when *B* is 9, 12, or 15, respectively. As expected, as the concentration of free P increases, and therefore *A* increases, the occupancy for the large *N* limit, *s*, also increases. The slope of *s* near *A* = 0 is steeper for larger *B* because the occupancy is more cooperative. Fig.2(D), (E), and (F) shows how the correlation length *ξ*_*c*_ depends on *A*. The correlation length *ξ*_*c*_ near *A* = 0 is also greater for larger *B* as well. *B* of 9, 12, or 15 correspond to interaction energy of about 5.4, 7.2, or 9.0 kcal mol^*−*1^, respectively, for the temperature of 300 K or 27°C.

**Fig. 2.**
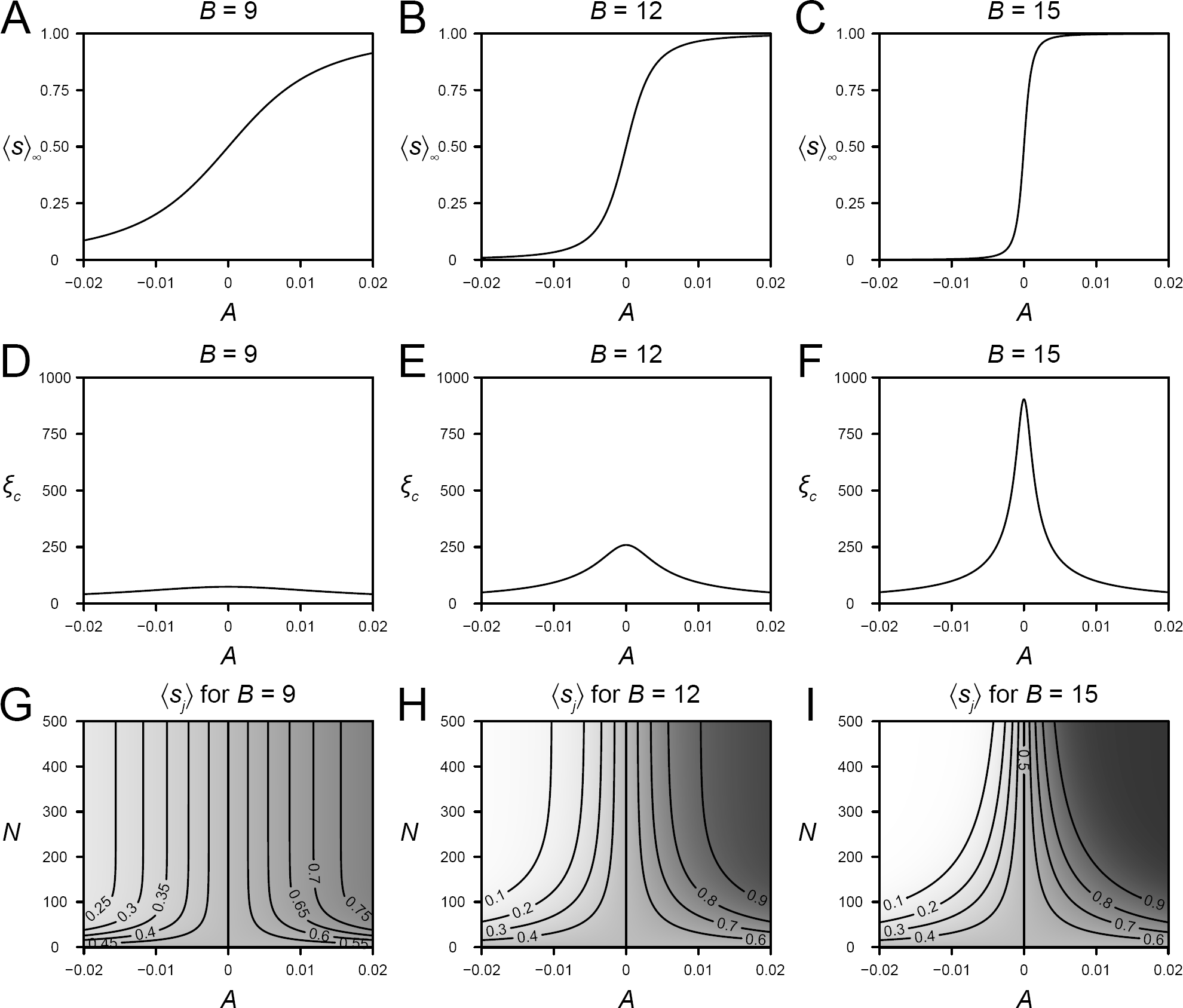
(A), (B), and (C) The relationship between ⟨*s*⟩ _*∞*_ and *A* = ln [*P*]_0_ + const. for *B* = 9, *B=* 12, and *B* = 15. (D), (E), and (F) The relationship between *ξ*_*c*_ and *A=* ln [*P*]_0_ + const. for *B=* 9, *B=* 12, and *B=* 15. (G), (H), and (I) Heat maps showing how the occupancy ⟨*s*_*j*_⟩ depends on *A* = ln [*P*]_0_ + const. and the telomere size *N* for *B* = 9, *B* = 12, and *B* = 15. When the cooperativity is weak (*B* is small), the length scale *ξ*_*c*_ remains small and the occupancy ⟨*s*_*j*_⟩ rapidly approaches ⟨*s*⟩_*∞*_ as the function of the telomere size *N*. However, when the cooperativity is strong (*B* is large), the length scale *ξ*_*c*_ can become large and the occupancy ⟨*s*_*j*_⟩ approaches ⟨*s*⟩_*∞*_ more slowly as the function of the telomere size *N*. When the abundance of the protein is high (*A* is large), the same occupancy level ⟨*s*_*j*_⟩ is achieved with smaller telomere size *N*.

Fig.2(G), (H), and (I) show how the general occupancy ⟨*s*_*j*_⟩ depends on *N* as well as as well as *A* = ln [*P*]_0_ + const. When the correlation length *ξ*_*c*_ is small, *s*_*j*_ rapidly approaches *s*. However, when *B* is large, there is a regime where *s* is close to 1, but *ξ*_*c*_ is large. In such a regime, the telomere achieves near full occupancy if *N* is large, but it will have a fair portion of unoccupied sites if *N* is smaller than *ξ*_*c*_. Therefore, the degree to which the telomerase activity is inhibited by P will depend on the size of the telomere *N* in this regime. Efficient inhibition of the telomerase activity is only achieved if *N* exceeds *ξ*_*c*_.

Now let us consider how the telomerase activity will depend on the abundance of the protein P. If the abundance is low and *A* = (ln [*P*]_0_ + const.) is low, there will be insufficient binding of the protein P on the telomere and less inhibition of the telomerase activity. This corresponds to the case when expression of the telomere binding protein is downregulated. Since there is less inhibition of the telomerase activity, the telomere length increases. If, on the other hand, the abundance of P is high, and therefore *A* is large, there will be near full occupancy even for low *N*. Thus the telomerase activity will be inhibited even for a short telomere. This leads to shortening of the telomere because of incomplete replication.

Our motivation was to come up with a mechanism of telomerase inhibition that is dependent on the size of the telomere and the abundance of the telomere binding protein(s). Our simple model provides the desired properties: effective inhibition does not take place unless the telomere size *N* exceeds the characteristic length *ξ*_*c*_; if the abundance of the protein is high, the telomerase activity is inhibited at a shorter telomere size *N*.

## Discussion

In this study, we presented a simple model of telomere length regulation. The model described binding of the inhibitory protein P to the telomere. As we have seen, by simply modeling the T-loop as a ring, and by simply postulating that binding of P is cooperative, the average occupancy of P per site can be sensitive to the size of the ring for some suitable ranges of the parameters. The protein P achieves near full occupancy only if the size of the telomere exceeds the correlation length. Therefore, the degree to which the telomerase activity is inhibited depends on the size of the telomere.

It is crucial that the telomere forms a T-loop, and therefore it does not have open ends. The presence of open ends would destabilize the binding of P near the ends. Thus, effective inhibition of telomerase activity requires T-loop formation.

The parameters need to be in an appropriate range for this mechanism to work. The cooperativity of the protein binding, represented by *B* = *b/* (*k*_B_*T*), needs to be strong enough. It may also seem as if the mechanism only works for a narrow range of the free protein concentration [*P*]_0_. However, the total number of the protein molecules is the sum of the numbers of the free and bound protein molecules. A narrow range of free protein molecules can correspond to a broader range of total protein molecules. Thus, the mechanism does not need be as sensitive to the abundance of the total protein.

Teixeira et al. (11) found that telomeres switch between two states: one that allows elongation by the telomerase and one that prohibits it. The mechanism based on cooperativity of the protein binding fits with such a switch-like phenomenon.

The model presented here is admittedly a simple toy model. A crucial question is whether the picture provided by this model is close to the actual mechanism. The real telomeres are more complicated because many different proteins bind to the telomeres and they vary depending on the organisms. The proteins likely form complex multivalent protein-protein interactions (21) and they will not be the simple nearest-neighbor interactions depicted in the model here. Nevertheless, what is essential to the mechanism proposed here is not the particular interaction presented in the model, but the cooperativity of the protein binding. It is possible that the multivalent interactions of the actual proteins stabilize their occupancies in a way that make them cooperative. If the cooperativity of the protein binding is strong enough, there can be length-dependence.

The model suggests that occupancies of some, if not all, telomere-binding proteins are dependent on telomere length. It would be interesting to study various telomere-binding proteins to see if any of them occupy telomeres in a length-dependent way.

## Acknowledgments

I would like to thank Dr. Bing Ren and Dr. David Gorkin for their critical reading of the manuscript.

## References

1. Blackburn EH, Greider CW, Szostak JW (2006) Telomeres and telomerase: the path from maize, tetrahymena and yeast to human cancer and aging. Nat. Med. 12:1133–1138. https://doi.org/10.1038/nm1006-1133

2. Greider CW, Blackburn EH (1985) Identification of a specific telomere terminal transferase activity in tetrahymena extracts. Cell 43:405–413. https://doi.org/10.1016/0092-8674(85)90170-9

3. de Lange T (2009) How telomeres solve the end-protection problem. Science 326:948–952. https://doi.org/10.1126/science.1170633

4. Griffith JD, et al. (1999) Mammalian telomeres end in a large duplex loop. Cell 97:503–514. https://doi.org/10.1016/S0092-8674(00)80760-6

5. Bianchi A, Shore D (2008) How telomerase reaches its end: mechanism of telomerase regulation by the telomeric complex. Mol. Cell 31:153–165. https://doi.org/10.1016/j.molcel.2008.06.013

6. Palm W, de Lange T (2008) How shelterin protects mammalian telomeres. Annu. Rev. Genet. 42:301–334. https://doi.org/10.1146/annurev.genet.41.110306.130350

7. Pfeiffer V, Lingner J (2013) Replication of telomeres and the regulation of telomerase. Cold Spring Harb. Perspect. Biol. 5:a010405. https://doi.org/10.1101/cshperspect.a010405

8. Nandakumar J, Cech TR (2013) Finding the end: recruitment of telomerase to telomeres. Nat. Rev. Mol. Cell Biol. 14:69–82. https://doi.org/10.1038/nrm3505

9. Murray AW, Claus TE, Szostak JW (1988) Characterization of two telomeric DNA processing reactions in saccharomyces cerevisiae. Mol. Cell. Biol. 8:4642–4650. https://doi.org/10.1128/MCB.8.11.4642

10. Marcand S, Brevet V, Gilson E (1999) Progressive cis-inhibition of telomerase upon telomere elongation. EMBO J. 18:3509–3519. https://doi.org/10.1093/emboj/18.12.3509

11. Teixeira MT, Arneric M, Sperisen P, Lingner J (2004) Telomere length homeostasis is achieved via a switch between telomerase- extendible and -nonextendible states. Cell 117:323–335. https://doi.org/10.1016/S0092-8674(04)00334-4

12. Marcand S, Gilson E, Shore D (1997) A protein-counting mechanism for telomere length regulation in yeast. Science 275:986–990. https://doi.org/10.1126/science.275.5302.986

13. Rodriguez-Brenes IA, Peskin CS (2010) Quantitative theory of telomere length regulation and cellular senescence. Proc. Natl. Acad. Sci. U. S. A. 107:5387–5392. https://doi.org/10.1073/pnas.0914502107

14. Greider CW (2016) Regulating telomere length from the inside out: the replication fork model. Genes Dev. 30:1483–1491. https://doi.org/10.1101/gad.280578.116

15. Hodges C, Crabtree GR (2012) Dynamics of inherently bounded histone modification domains. Proc. Natl. Acad. Sci. U. S. A. 109:13296–13301. https://doi.org/10.1073/pnas.1211172109

16. Metropolis N, Rosenbluth AW, Rosenbluth MN, Teller AH, Teller E (1953) Equation of state calculations by fast computing machines. J. Chem. Phys. 21:1087–1092. https://doi.org/10.1063/1.1699114

17. Glauber RJ (1963) Time-Dependent statistics of the ising model. J. Math. Phys. 4:294–307. https://doi.org/10.1063/1.1703954

18. Ising E (1925) Beitrag zur theorie des ferromagnetismus. Zeitschrift für Physik 31:253–258. https://doi.org/10.1007/BF02980577

19. Ishii H (2000) A statistical-mechanical model for regulation of long-range chromatin structure and gene expression. J. Theor. Biol. 203:215–228. https://doi.org/10.1006/jtbi.2000.1081

20. Kramers HA, Wannier GH (1941) Statistics of the Two-Dimensional ferromagnet. part I. Phys. Rev. 60:252–262. https://doi.org/10.1103/PhysRev.60.252

21. Shi T, et al. (2013) Rif1 and rif2 shape telomere function and architecture through multivalent rap1 interactions. Cell 153:1340–1353. https://doi.org/10.1016/j.cell.2013.05.007

